# The representation of voluntary and reflexive fast eye movements in the macaque lateral intraparietal area (LIP)

**DOI:** 10.64898/2026.02.10.705045

**Authors:** Sadra Fathkhani, Bahareh Taghizadeh-Sarshouri, Andre Kaminiarz, Frank Bremmer

**Author notes:** Corresponding Autor: Sadra Fathkhani. Shared first-authorship (Sadra Fathkhani, Bahareh Taghizadeh-Sarshouri).

## Abstract

The macaque lateral intraparietal area (LIP) is known for its role in visually guided saccades as well as in higher cognitive functions. However, its contribution to more basic visuomotor processes remains unclear. Here, we investigated whether neural activity in area LIP is also related to involuntary reflexive eye movements, specifically the fast phases of optokinetic nystagmus (OKN). To address this question, we compared spiking activity and local field potentials (LFPs) in area LIP of two male macaque monkeys during visually guided saccades and during kinematically similar OKN fast phases. Neurons exhibiting robust perisaccadic activation during voluntary saccades showed markedly reduced or no activity around the time of OKN fast phases and were not modulated by fast-phase amplitude or frequency. Using a Generalized Linear Model, we found that during OKN slow phases, area LIP reliably encoded gaze position and the direction of visual motion driving the reflexive eye movement. LFP analyses further revealed that beta-band power differed between voluntary saccades and OKN fast phases, whereas theta-band phase coherence increased following both types of fast eye movements, suggesting distinct local processing but shared post-movement network coordination. Our results reinforce the view of area LIP as a key area for integrating sensory and cognitive signals relevant for goal-directed action rather than a generic oculomotor controller.

## Introduction

The lateral intraparietal area (area LIP) in the macaque monkey is known for its involvement in the control of voluntary saccadic eye movements. Many neurons show spatially tuned activity for visually guided saccades (1–3), as well as modulation related to the direction of pursuit eye movements (4–7) and, to a lesser extent, to gaze velocity (8,9). Area LIP has also been implicated in higher cognitive functions such as salience, priority and attention. In these studies, neurons were shown to track the evolving attentional priority of locations, and respond most strongly to behaviorally relevant stimuli, suggesting the encoding of a salience map for flexible attentional selection (10,11). Likewise, it was shown that LIP neurons encode the intention to make a saccade, independent of immediate sensory or motor events (12,13) and to keep this information in spatial working memory (14). Remarkably, also other cognitive aspects are represented in area LIP. As an example, Janssen and Shadlen showed neural activity to reflect an internal representation of elapsed time (15). In another study monkeys were placed in a virtual dynamic foraging environment in which they had to track the changing values of alternative choices through time. Neurons were found to represent the relative value of competing actions (16). Even numerosity is represented in area LIP: Roitman and colleagues showed that a population of neurons in area LIP encodes the total number of elements within their classical receptive fields within the first 100 ms after stimulus onset in a graded fashion, across a wide range of numerical values, and that this encoding was independent of attention, reward expectations, or stimulus attributes such as size, density, or color (17). In summary, the contribution of area LIP to various cognitive processes has been widely studied and convincingly demonstrated. This is in stark contrast to our sparse understanding of area LIP’s basic visuo-motor functions.

In all the above-mentioned studies, monkeys were required to make voluntary, goal-directed saccades. A more complete understanding of the role of area LIP in oculomotor control, however, requires probing its neural activity also during involuntary eye movements. Such eye movements typically occur during oculomotor reflexes, including the optokinetic nystagmus (OKN) and the vestibulo-ocular reflex (VOR. (18)). OKN can be elicited simply by moving a cloud of random dots in front of an observer, causing the eyes to smoothly follow the pattern movement (slow-phase) interrupted with rapid resetting movements (fast-phases) in the opposite direction. The amplitudes of these involuntary fast eye movements vary and are typically larger during look-as compared to stare-nystagmus (19), the latter usually being characterized as more reflexive in nature. Moreoverover, the frequency of fast phases is higher during stare-as compared to look nystagmus. In humans, the so-called main-sequence of visually guided saccades and fast-phases of equal amplitude are similar but not identical: for a given amplitude, peak velocity is higher, and duration is shorter during voluntary saccades as compared to fast-phases (20). Early functional magnetic resonance imaging (fMRI) studies in human observers confirmed the role of parietal cortex in encoding the intention to make a saccade (21) and reported reduced Blood-oxygenation-level-dependent (BOLD) contrast in parietal cortex during stare-as compared to look-nystagmus (22).

Here we investigated the differential role of area LIP of the macaque monkey for voluntary and reflexive fast eye movements. More specifically, we compared spiking activity and Local Field Potentials (LFPs) in area LIP of two macaque monkeys during voluntary, goal directed saccades, and involuntary, reflexive fast-phases of OKN of similar metric (direction and amplitude). LFPs reflect the summated synaptic inputs and local processing within a neuronal population, whereas spiking activity primarily represents the output of individual neurons (23–25). We hypothesized reduced spiking output during involuntary, reflexive fast phases of OKN as compared to voluntary saccades of the similar metric, likely reflecting differences in the underlying input signals as indicated by the LFPs.

## Methods

### Animal

Behavioral and neural recordings were performed in two adult male rhesus macaques (Macaca mulatta). In two separate surgeries each monkey was implanted with a head holder and a recording chamber. Based on MRI scans acquired prior to surgery, the chamber (inner diameter 14 mm) was centered at a position 3 mm posterior from the inter-aural line and 15 mm lateral from the longitudinal fissure (P3/L15) in monkey S and at P5/L12.5 in monkey E to access the region of the IPS relevant for our recordings (right/left hemisphere in monkey S/E). During the experiments, the correct position of the electrode within area LIP was determined based on the functional properties of the neurons under study which was saccade direction selectivity. All procedures had been approved by the regional authorities and were in accordance with the published guidelines on the use of animals in research (European Communities Council Directive 2010/63/EU).

### Experimental set-up

Monkeys were sitting head-fixed in a primate chair in a completely dark room, and their eye position was monitored at 1000 Hz using a video-based eye tracker (EyeLink 1000, SR Research, Ottawa, Canada). The chair was placed 97 cm away from a semi-transparent screen (size 160 cm × 90 cm, subtending the central 79° × 50° of the visual field) on which the visual stimuli were back-projected using a PROPixx-projector (VPixx Technologies, St-Bruno de Montarville, Canada) running at a resolution of 1920 × 1080 pixels and at a frame rate of 100 Hz.

### Recording of neural activity

Spiking activity and local field potentials (LFP) were recorded using standard tungsten microelectrodes (FHC, Bowdoin, USA) with an impedance of ∼ 2 MΩ at 1 kHz that were positioned by a hydraulic micromanipulator (MO-95, Narishige, Tokyo, Japan). A stainless-steel guiding tube was used for transdural penetration and support of the electrode. The neuronal signal was recorded and amplified using a commercial system (Alpha Omega, Nof HaGalil, Israel), sampled at 44 kHz.

### Behavioral paradigms

#### Overlap saccade

Temporal and spatial parameters of this paradigm were adjusted so that the performance of each monkey was in the range of 90 %. Each trial began with presentation of a central fixation point (Figure 1a), i.e., a red square with a side length of 0.8°. Upon successful fixation within a predefined window (+/-2.5° by +/-2.5° monkey S, and +/-4° by +/-4° monkey E, centered on the fixation point) for a random duration (500-700 ms for monkey S, 600-1000 ms for monkey E), a peripheral saccade target (with the same physical specification as the central fixation) appeared at one of four possible locations on the cardinal axes at 10° eccentricity. The peripheral saccade stimulus remained visible on the screen for a random duration, 400-600 ms for monkey S and 50 ms for monkey E, while the monkey maintained central fixation. The central fixation point then disappeared (go signal), cueing the monkey to make a saccade towards the peripheral target within 1000 ms. Following the saccade and successful fixation of the peripheral target (600 ms for monkey S, and 300 ms for monkey E), liquid reward was delivered. Error trials were defined when the monkey did not initiate the saccade within 1000 ms after the go signal, or the saccade was made outside of an +/-4° by +/-4° window centered on the target. In order to compare neural activity along with goal-directed saccades and Optokinetic Nystagmus fast-phases, the preferred saccade direction (PD) of every neuron was estimated visually and auditorily during the experiment. This estimation was then used to determine the proper stimulus direction in the Optokinetic Nystagmus task.

**Figure 1.**
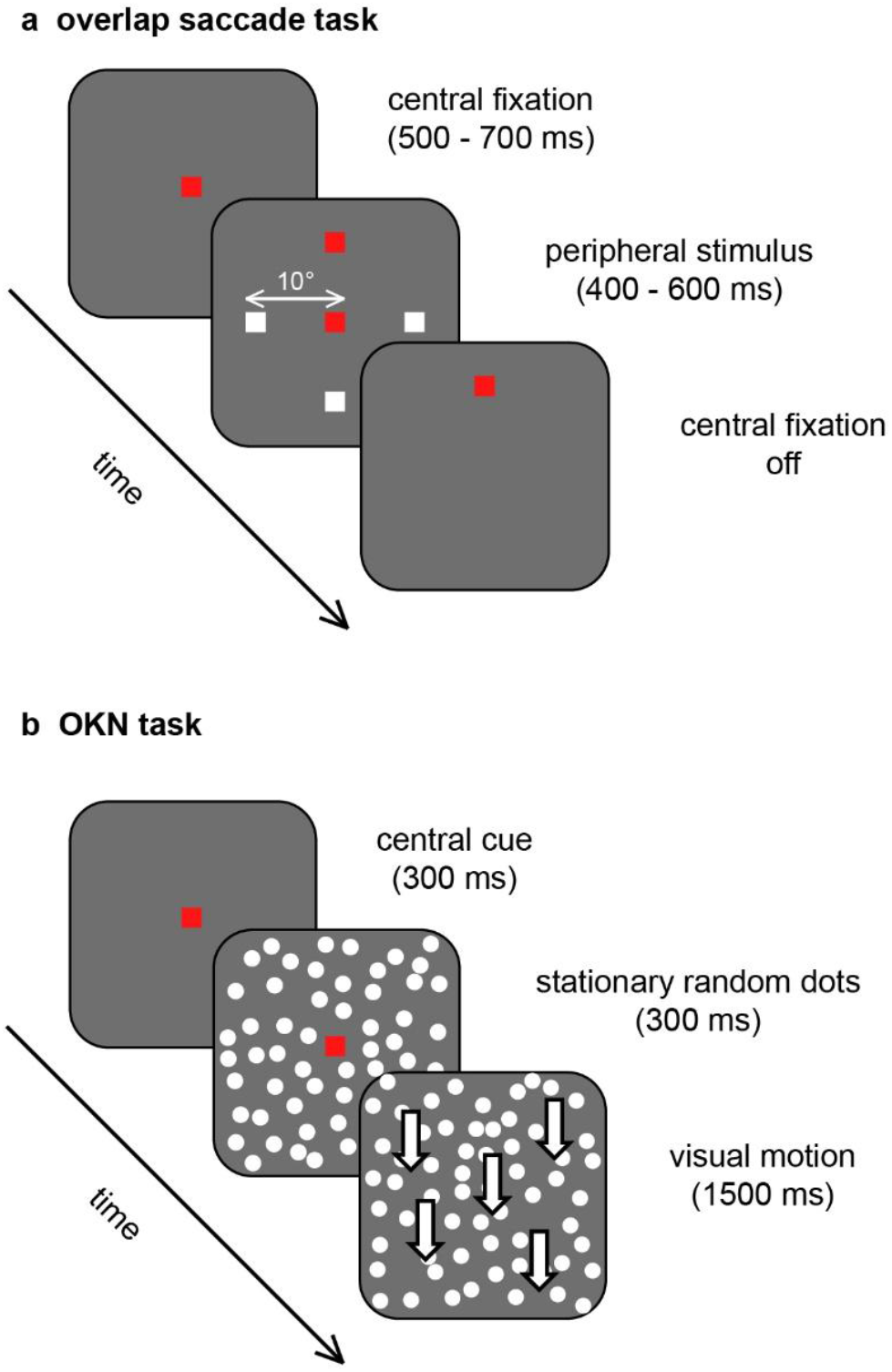
Behavioural paradigms. a. Visually guided overlap saccade task. Monkeys fixated a central red target. A peripheral red target (10° eccentricity) appeared at one of the four cardinal locations while fixation remained visible (90° in this example; the other potential target locations are shown in white only for illustrative purpose and were not visible to the monkey). After a random fixation-target overlap time, fixation offset served as the go cue, and the monkey made a saccade to the still visible target to obtain a liquid reward. b. Optokinetic Nystagmus (OKN). A central red square was presented as a starting cue, followed by a stationary random-dot pattern while the central cue remained visible. After central cue offset, all the dots moved in a cardinal direction (downward in this example), opposite to the preferred saccade direction of a neuron under study, eliciting OKN with fast-phases in the neuron’s preferred saccade direction.

#### Optokinetic Nystagmus (OKN)

OKN fast-phases in the PD of the neuron under study were induced by large field coherent visual motion in the direction opposite to the estimated PD (Figure 1b). Each trial began with the presentation of a central cue (a red square with a side length of 0.8°) on a black background, signaling the monkey that he could start the trial. The trial was initiated when the monkey’s gaze entered a +/-20° by +/-20° control window around the central cue. After 300 ms looking within this window, a Random Dot Pattern (RDP) containing 1000 white dots (with a diameter of 0.4 degrees each on a black background spanning a visual field of 55° horizontally and 45° vertically) was displayed for 300 ms (the stationary period). Then, the central cue disappeared, and the RDP started to move at 18.43 °/s or 16.46 °/s respectively in Y or X direction (motion period). Each element of the moving RDP had a limited lifetime of 10 frames (=100 ms) to prevent the monkeys from tracking individual dots throughout the trial. Following the motion period, the RDP was switched off, and liquid reward was delivered to the monkeys. Error trials were defined when the monkey looked outside of the +/-20° by +/-20° window during the motion period. In 16 sessions from monkey S and 14 sessions from monkey E additional trials were recorded where the visual motion was presented in the preferred saccade direction of the neuron under study to evoke fast-phases in the opposite direction. In case of two motion directions, these were presented randomly interleaved across trials.

### Behavioral data analysis

Saccades and fast-phases were identified offline with a custom written script (Matlab, The Mathworks Inc., Ithaca, USA). The raw eye-position signal was convolved with a 5-ms Gaussian kernel, and onset was defined when eye velocity exceeded 80°/s for three consecutive samples. Offset was determined when velocity remained below this threshold for three consecutive samples. To avoid contamination by non-directional LIP transient activity (26) and mixed smooth eye movement responses (27,28), fast phases within 500 ms after stimulus motion onset were excluded. The main-sequence computation was used to analyze the dynamics of fast eye movements, describing the relationship between the maximum velocity and duration of fast eye movements with their amplitudes (29). For each monkey the main-sequences of goal-directed saccades and fast-phases within an amplitude range of 2-10 degrees were separately fitted to a power function (eq. 1) using a non-linear least-square fitting method (30). The model parameters (*a* and *b*) for the main-sequence regarding maximum velocity and duration were compared between saccades and fast-phases using a two-sample z-test.

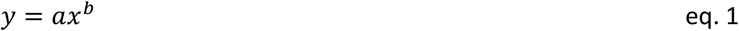

### Neural data analysis

#### Spiking activity

Spike-sorting was performed offline using the Plexon Offline Sorter V3.3.5 (Plexon Inc. Dallas, United States). Quantitative directional tuning was determined from the overlap saccade task. For each neuron, the trial-averaged firing rate (see below) was computed from a 100 ms window around saccade onset, that is, within −50 to 50 ms of saccade onset for monkey S and 0 to 100 ms for monkey E. Different time windows were used for the two monkeys to center the window on the respective time of the population peak response. Neurons showing significant direction-tuning (one-way ANOVA on neural activity with saccade direction as factor, p < 0.05) and which also had been recorded during OKN were included in further analyses. Preferred saccade direction was defined as the direction with the highest response within the specified analysis window.

For each trial (OKN or saccade), a spike density function (SPD) was computed at 1-ms resolution using the *psth()* function from the Chronux toolbox (31), with a 25-ms Gaussian kernel. For a given OKN trial, the SPD was analyzed in a 300 ms time window spanning −150 to +150 ms relative to fast-phase onset for monkey S and −50 to +250 ms for monkey E. This procedure was repeated across all trials, and the resulting SPDs were averaged to determine a neuron’s mean response for a given fast-phase direction. For goal directed saccades, SPDs were analyzed in the same way, using the same temporal windows (−150 to +150 ms for monkey S and −50 to +250 ms for monkey E), relative to saccade onset. The average SPD for each saccade direction was used in subsequent analyses. For both animals, pre-movement activity was defined as the mean firing rate during the 50 ms period preceding eye movement onset (−50 to 0 ms). Post-movement activity was assessed between 25–75 ms for monkey S and 50–100 ms for monkey E relative to eye movement onset.

Optokinetic nystagmus typically comes at one of two flavors: either stare- or look-nystagmus (22,32). Both are characterized by the frequency and amplitude of their fast phases: around 3 Hz and with smaller amplitudes for stare-nystagmus and around 1 Hz and with larger amplitudes for look-nystagmus. Here, we determined the frequency of fast-phases as the inverse of the temporal distance between the onset of two consecutive fast-phases. To analyze the effect of fast-phase frequency and amplitude on neural activity, for every neuron, fast-phases with an amplitude between 2 and 10 degrees that occurred at frequencies between 1 and 4 Hz were grouped into bins with the lower edge of each bin included.

Only saccades and fast-phases with comparable spatial (direction and amplitude) and kinematic properties were analyzed for their relationship with spiking and LFP activities. To this end, fast-phases had to match the following criteria: (1) amplitude between 5° and 10° (the average saccade amplitude in the overlap task was 7°, see Figure 2), (2) direction within 45° of the saccade PD, and (3) a minimum interval of 150 ms between two consecutive fast-phases.

**Figure 2.**
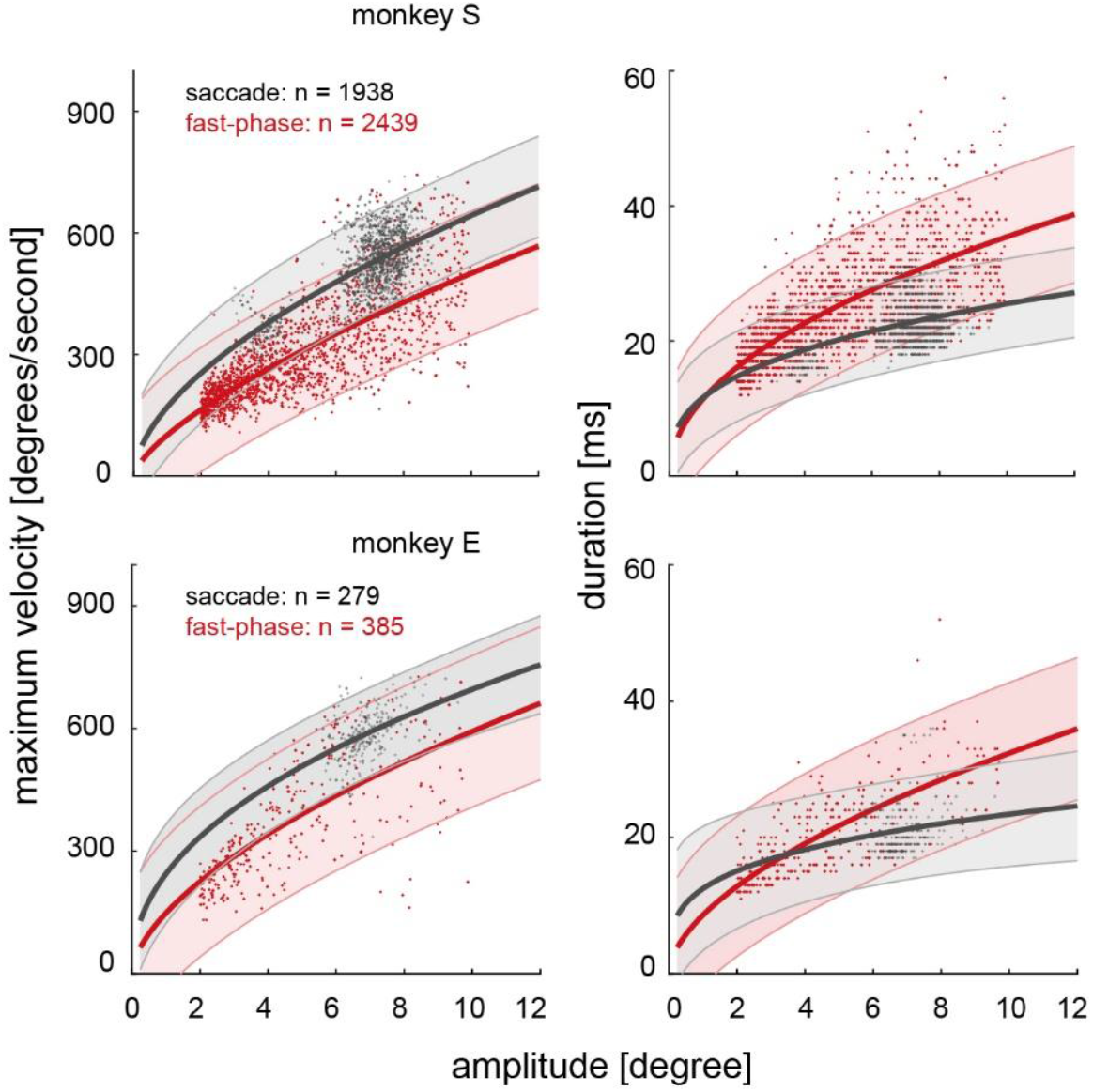
Main-sequence of saccades and fast-phases. Each row shows data from one animal. Left and right columns illustrate peak velocity and duration, respectively, as a function of amplitude. Every point represents one saccade (black) or fast-phase (red). Thick solid curves illustrate power-function fits computed separately for saccades and fast-phases. Shaded areas indicate the corresponding 95% confidence intervals of the fits.

### Generalized Linear Model (GLM) of the spiking activity during the slow-phase

For each neuron, we extracted SPDs around all fast phases fulfilling the following criteria: amplitude range of 2°–10°; inter-fast-phase interval ≥ 150 ms. For each fast phase, we analyzed two time windows separately: the 100 ms preceding fast-phase onset (corresponding to the end of the slow phase) and the 100 ms following fast-phase offset (the beginning of the next slow phase). In each window, we modelled neural activity using a generalized linear model (GLM) that included horizontal and vertical gaze position (*g*_*x*_, *g*_*y*_), horizontal and vertical gaze velocity (*v*_*x*_, *v*_*y*_), and the direction of stimulus motion (*MoDir*) relative to the neuron’s preferred direction as predictors.

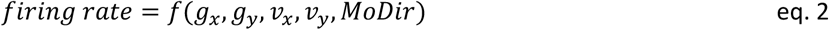

The factor *MoDir* was assigned a value of 1 for trials in which the stimulus moved in the same direction as neuron’s saccade PD, and, −1 when the stimulus moved in the opposite direction. All recorded neurons were included in this analysis, regardless of whether or not they exhibited a significant saccade tuning. Model parameters were standardized, and we used a normal distribution and identity link function for the GLM. The model was fitted using the SPD and the corresponding predictor values across all time points within the respective time intervals and all detected fast phases. Models were implemented using the *glmfit()* function in MATLAB. The significance of the coefficients was determined using the reported p-values by the function.

### Pre-processing of local field potentials

the raw LFPs were measured with 1 KHz temporal sampling rate. The LPF trace from every session was notch filtered at 50 Hz, low pass filtered at 100 Hz, then sorted into trials from 500 ms before the trial initiation to 500 ms after reward time. For every trial the LFP power spectrum was calculated in 1 Hz frequency bands within the 2.5-40 Hz frequency range using the MATLAB Morse wavelet transform (*cwt()* function). LFP phase angle and power were calculated as the angle and amplitude of the wavelet transform output. The power in each frequency band was normalized relative to the trial’s baseline within the same frequency band using the following equation:

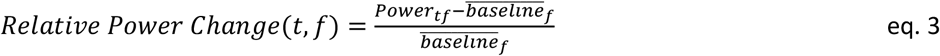

where *Power*_*tf*_ is the power at time *t* and frequency *f*, and 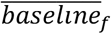 is the power in frequency *f* during the 300 ms interval before onset of the stationary dot pattern in OKN trials. In the saccade task the last 300 ms of the fixation period before the step of the saccade target was taken as baseline. Normalization relative to the frequency-specific baseline accounts for both trial-by-trial variability and 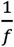 power distribution (33). Inter-trial Phase Clustering (ITPC) or intertrial phase coherence at time *t* and frequency *f* quantifies the uniformity of the distribution of the phase angles across trials at each time-frequency point in the polar space (33). It represents the magnitude of the average vector computed from all phase vectors, where every phase vector is defined by a unit length and the LFP phase angle at time *t* and frequency *f*. In all reported figures and results where spiking activity, LFP power and phase were analyzed before and after a fast-phase/saccade, the signals were aligned once to the onset and once to the offset of the fast-phase/saccade.

## Results

Throughout the 55 recorded sessions (38 for monkey S and 17 for monkey E) a total of 8638 OKN fast-phases (7542 for monkey S, 1096 for monkey E) were detected. 2939 of those occurred within the first 500 ms after motion onset and were excluded (see Methods), leaving 5699 (4942 for monkey S, 757 for monkey E) fast-phases for further analysis. Likewise, we recorded 1938 saccades in monkey S and 279 saccades in monkey E.

### The main-sequence

We compared eye movement dynamics between saccades and OKN fast-phases using the main-sequence approach (see Methods). Figure 2 illustrates the dependence of maximum velocity (left column) and duration (right column) on movement amplitude ranging from 2° to 10°, for saccades (black) and OKN fast-phases (red) for each monkey (top row: monkey S; bottom row: monkey E). For both, maximum velocity and amplitude, respectively, the relationships were well described by a power-law model (20,30). The corresponding fit parameters are listed in Table 1. A two-sample z-test revealed that fast-phases exhibited significantly lower maximum velocity than saccades (Table 2), consistent with previous findings in human participants (20,34). Although, in both monkeys, fast-phases tended to be longer than saccades, differences were not statistically significant.

**Table 1.**
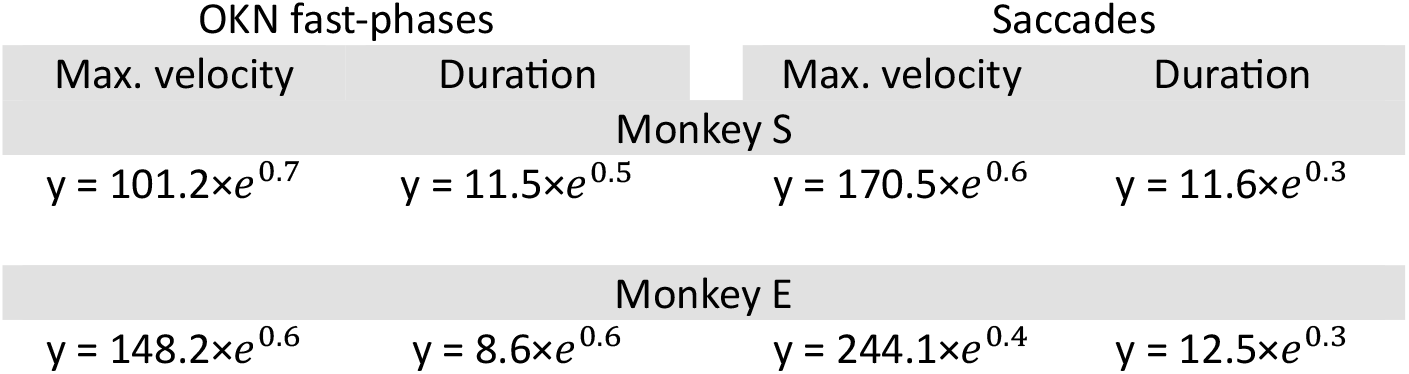
Estimated parameters of power function fits.

| OKN fast-phases |  | Saccades |  |
| --- | --- | --- | --- |
| Max. velocity | Duration | Max. velocity | Duration |
| Monkey S |  |  |  |
| $y = 101.2 \times e^{0.7}$ | $y = 11.5 \times e^{0.5}$ | $y = 170.5 \times e^{0.6}$ | $y = 11.6 \times e^{0.3}$ |
| Monkey E |  |  |  |
| $y = 148.2 \times e^{0.6}$ | $y = 8.6 \times e^{0.6}$ | $y = 244.1 \times e^{0.4}$ | $y = 12.5 \times e^{0.3}$ |

**Table 2.**
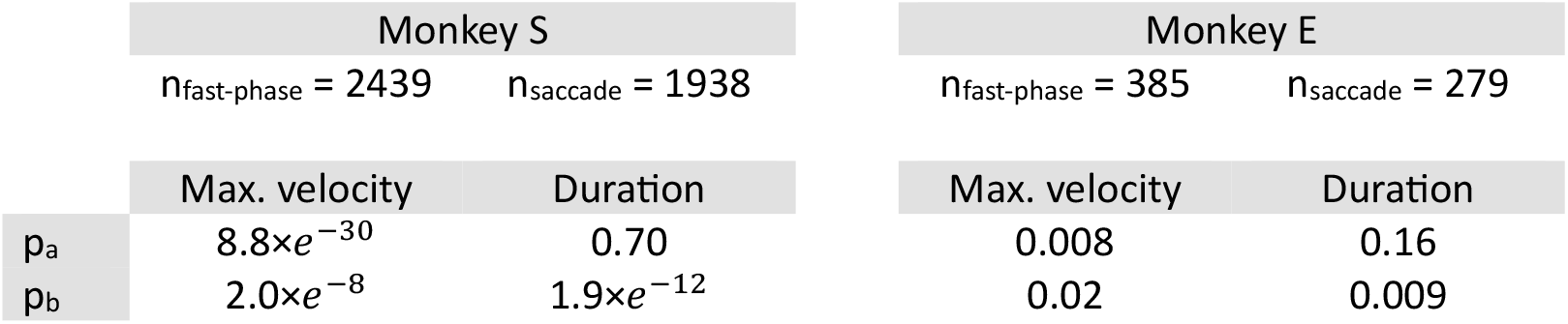
Two-sample z-test on the power function parameters between OKN fast-phases and saccades.

### Saccade and fast-phase related neural activity

Next, we examined whether neurons showing saccade-related activity were also active during fast phases. To this end, we first identified neurons that were tuned for saccades and then compared their activity around the onset of fast phases and saccades in the neuron’s preferred saccade direction. From the full sample of 135 neurons (98 in monkey S, 37 in monkey E), 70 neurons (59 in monkey S and 11 in monkey E) exhibited significant saccade direction tuning (see Methods). For 57 neurons (50 in monkey S and 7 in monkey E) the stimulus motion direction in the OKN task was correctly set opposite to the neuron’s preferred saccade direction to evoke fast-phases in the saccade PD; these neurons were included in further analyses. Neural activity was compared between kinematically similar saccades and fast-phases (see Methods). Figure 3a illustrates the average response of a representative neuron around the onset of saccades (black) and fast-phases (red) in the saccade PD. This neuron preferred downward saccades (270°; Figure 3b). Figure 3d demonstrates the average response of the same neuron to the saccade onset (black) and target onset (grey), indicating stronger selectivity for the saccade as compared to the visual stimulus. In the OKN task, upward (90°) dot motion elicited downward (270°) fast-phases. Figure 3c shows the average vertical eye position aligned to movement onset, corresponding to Figures 3a and 3b. Despite having similar kinematics, the neural activity shows drastically distinct patterns between the two movements. While neural activity started rising shortly before the saccade onset, it was not modulated around the onset of fast-phases. This reduction in spiking output was the typical distinction we observed across neurons. To extend the comparison to the population of neurons, trial-averaged neural activity around saccade and fast-phase onset were computed for each neuron’s saccade PD. Figure 4 shows the average neural firing rate around saccades (horizontal axis) and fast-phases (vertical axis) in a time window −150 to +150 ms relative to saccades/fast-phases onset for monkey S and −50 to +250 ms for monkey E (details in Methods). On average the activity was higher for saccades as compared to fast-phases (one-sided right-tailed Wilcoxon signed-rank test on the difference of activity between the tasks: n = 57, p < 0.001). We also compared neural activity before and after saccades/fast-phases (Figure 5). A Wilcoxon signed-rank test revealed a significant difference between pre- and post-saccadic spike count rates (monkey S: n = 50, p = 0.02; monkey E: n = 7, p = 0.01), whereas, no significant difference between pre- and post-fast-phase activity was found (monkey S, n = 50, p = 0.73; monkey E, n = 7, p = 0.2), indicating that modulations occurred specifically around saccades. We also compared pre- and post-movement activity directly between saccades and fast-phases. Post-saccadic and post-fast-phase spike count rates differed significantly in both animals (monkey S, p < 0.0001; monkey E, p = 0.03). Pre-saccade and pre-fast-phase activity also differed significantly in monkey S (p = 2.8 ×*e*^−9^), but showed only a trend in monkey E (p = 0.08). The difference between the two monkeys could be explained by the temporal difference in the peak of the population response: for monkey E the population response had a post-saccadic peak, while it was before saccade onset in monkey S.

**Figure 3.**
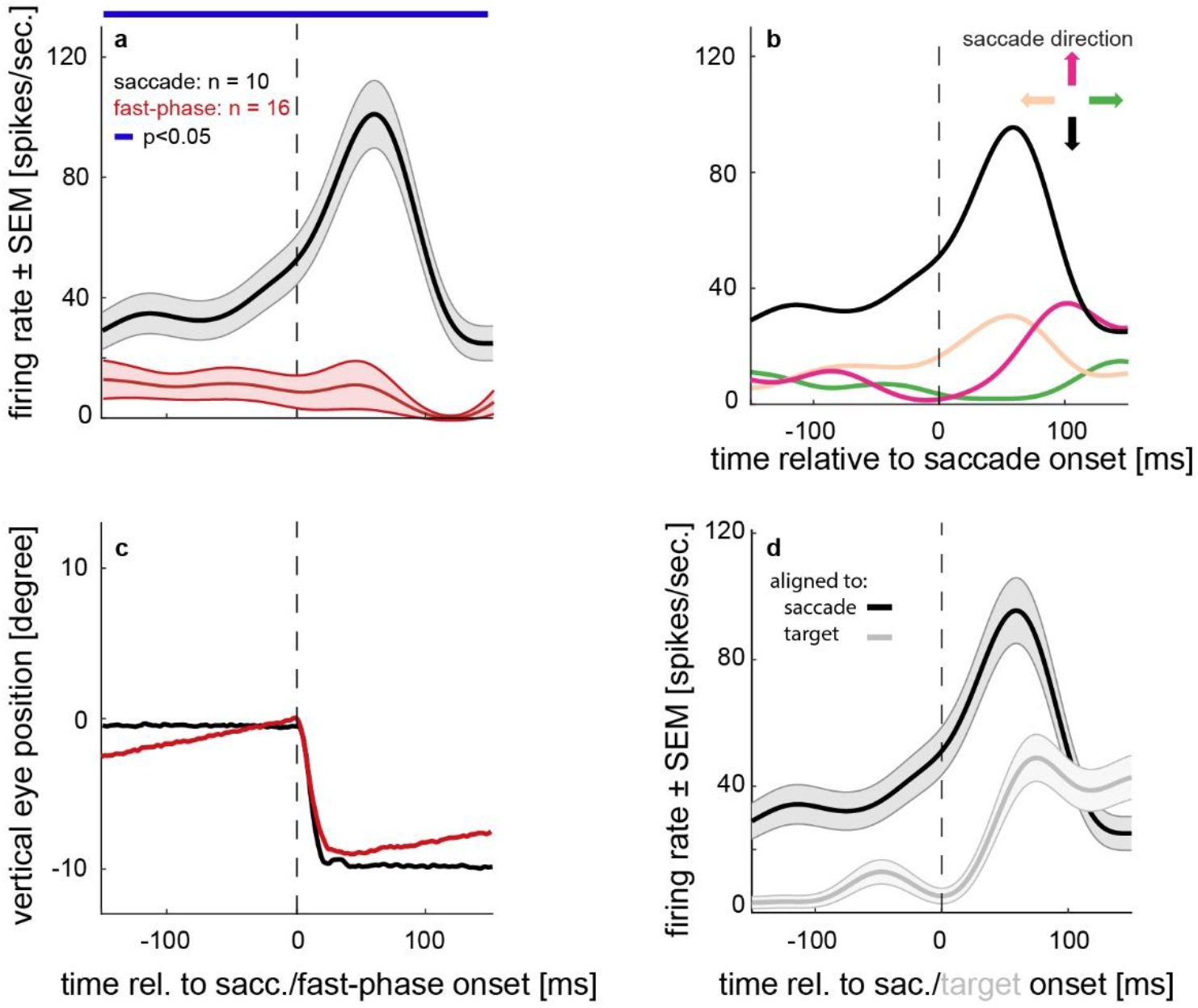
Average spike density function (SPD) of a representative neuron. **a** Neural activity around the onset of saccades (black) and OKN fast-phases (red). Shaded areas show the standard error of the mean (SEM). The inset indicates the number of trials, and the dashed vertical line marks the saccade/fast-phase onset time. The blue horizontal line shows the time bins with a significant difference between the two curves (see Methods). **b** Average activity of the same neuron when making saccades to four cardinal targets, where the color-coded arrows and lines indicate saccade target directions. **c** Average vertical eye position corresponding to **a**. Colour coding and other settings are similar to **a. d** Average activity of the same neuron aligned to saccade onset (black) and target onset (grey). Like before, shaded areas indicate the SEM.

**Figure 4.**
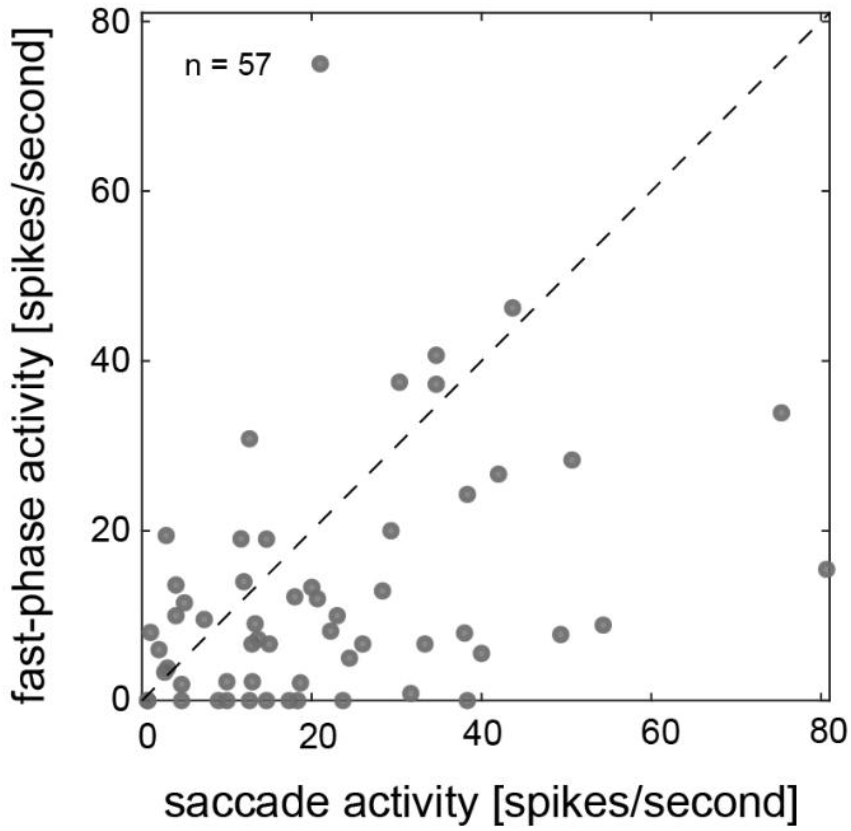
Trial average activity around the time of saccades and fast-phases. Neural activity around the time of saccades (horizontal axis) and OKN fast-phases (vertical axis), both in a given neuron’s preferred saccade direction. Each dot represents data from a given neuron.

**Figure 5.**
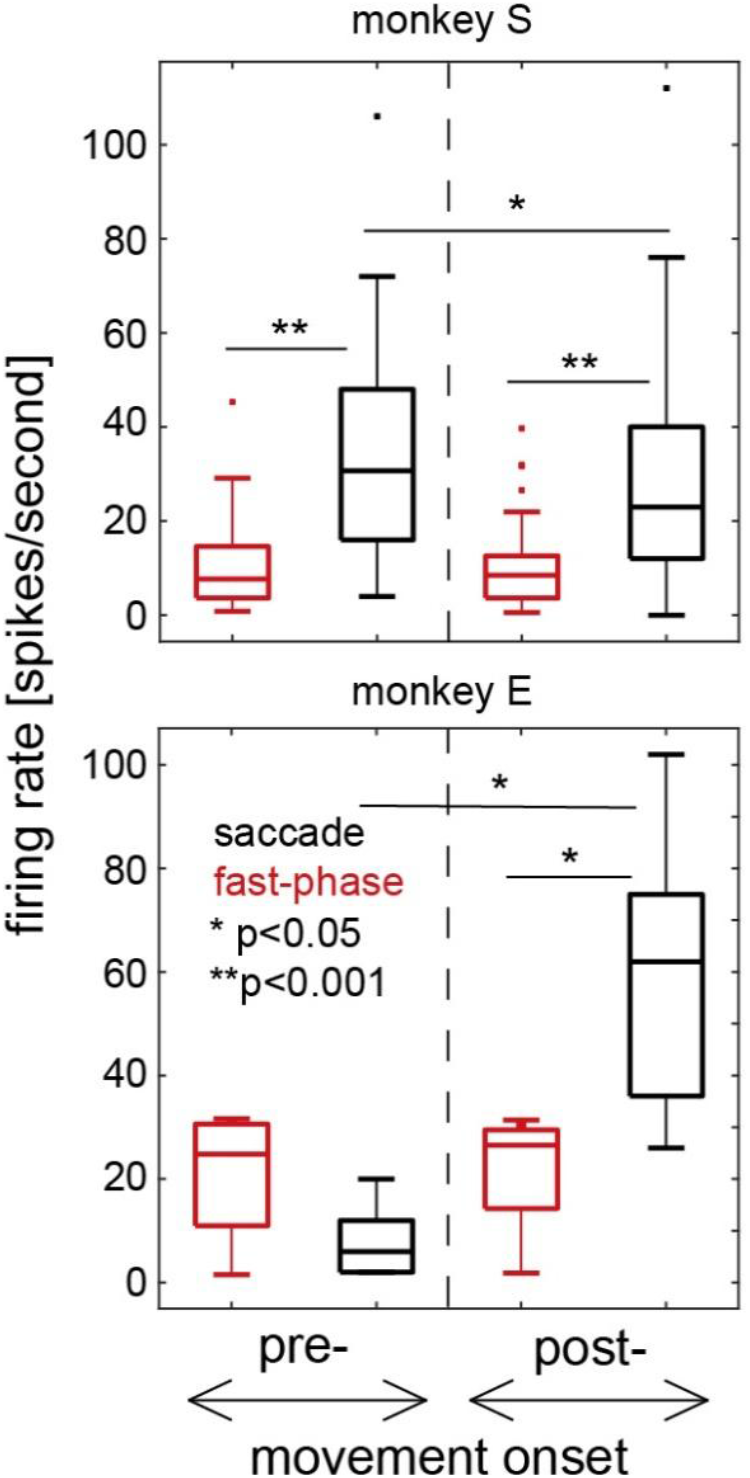
Average neural activity before and after saccade and fast-phase onset. The dashed vertical line separates neural activity before (left) and after (right) saccade (black)/fast-phase (red) onset. Every box plot shows the median and the 75th (top) and 25th (bottom) percentiles, whiskers indicate the data range within 1.5 times the interquartile range, and dots show data points beyond whiskers (outliers).. Data for monkey S are shown the top panel, data for monkey E in the bottom panel. Statistics (p-values) indicate the result of a one-sided (right-tailed) Wilcoxon signed-rank test on the difference of activity between the time intervals/tasks.

### Modulations of neural activity by the frequency and the amplitude of fast-phases

Functional imaging studies on human participants have reported that the frequency of fast-phase occurrences modulates BOLD contrasts (see Introduction). Specifically, increased fast-phases frequency, typical for stare nystagmus, is associated with decreased BOLD activity in cortical oculomotor centers, while lower-frequency fast-phases during look-nystagmus lead to a broader activation across parietal regions (22,35). Here we tested whether such systematic modulations are also reflected in neural activity in area LIP. To this end, for each neuron, fast-phases in saccade PD were sorted into 1-Hz frequency bins and neural activity was calculated around their onset. After amplitude and frequency filtering (see Methods), 1634 fast-phases from 50 neurons remained for further analysis from monkey S and 130 from 7 neurons for monkey E. Figure 6 (left column) illustrates the average spike rate of the population of tuned neurons as a function of fast-phase frequency. No significant effect of frequency on spike rates was observed (Kruskal-Wallis test; monkey S, p = 0.27; monkey E, p = 0.82). Additionally, we examined whether fast-phase amplitude modulates neural activity in area LIP. After amplitude filtering (see Methods), 2439 fast-phases from 50 neurons of monkey S and 385 from 7 neurons for monkey E were included in the analysis. Fast-phases were sorted into 1° amplitude bins, regardless of their frequency, and neural activity was again calculated around their onset. Figure 6 (right column) illustrates the average spike rate of saccade-tuned neurons as a function of fast-phase amplitude. No significant effect of amplitude on activity was observed across the full amplitude range (Kruskal-Wallis test: monkey S, p = 0.12; monkey E, p = 0.72). Taken together, these results suggest that neither fast-phase frequency nor amplitude modulate neural activity in area LIP.

**Figure 6.**
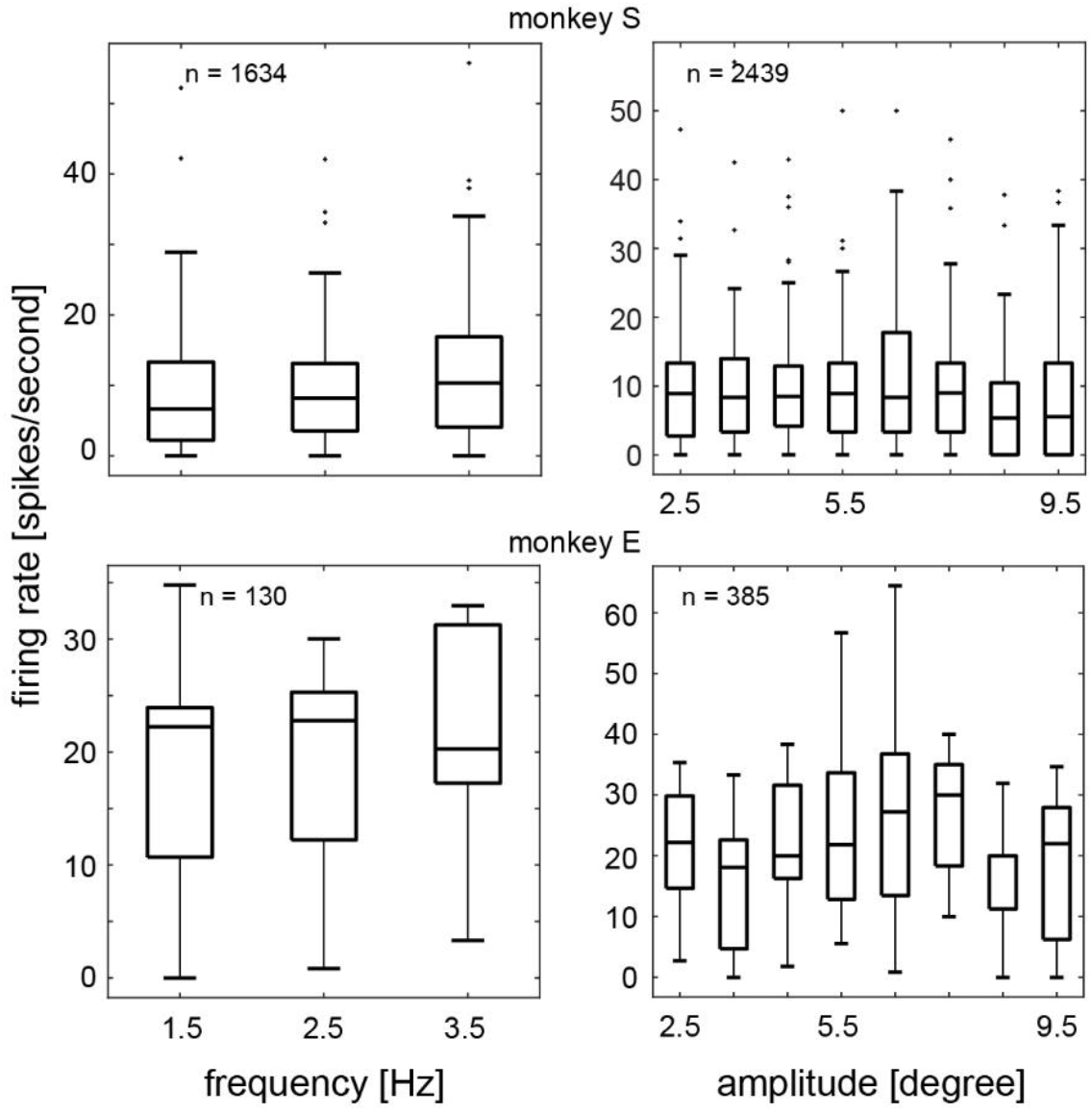
Neural activity as a function of frequency and amplitude of fast-phases. Every box plot shows the median and the 75th (top) and 25th (bottom) percentiles, whiskers indicate the data range within 1.5 times the interquartile range, and dots show data points beyond whiskers (outliers).. Number of fast-phases included in every analysis is indicated as inset (see Methods).

### Neural activity during the OKN slow-phase

The central goal of this study was to examine whether the modulatory effects of eye movement parameters in LIP differ between voluntary and involuntary movements. During goal directed behavior, spiking activity in LIP is modulated by eye position (7,36–38) and, to some extent, by eye velocity. While dominated by saccadic eye movements, neurons in LIP have also been reported to be selective to the direction of pursuit eye movements (7), as well as to the direction of the moving dots stimulus within their receptive field, even when the stimulus is not behaviorally relevant (4,39–43). Here we asked whether these modulations also occur during involuntary, non-goal directed behavior. To address this, we fitted GLMs to the firing rate of each neuron at the beginning and end of the slow-phase of the OKN. During these fast-phase-free epochs, the same factors —such as eye position, velocity, and stimulus direction— could potentially modulate the activity.

For a subset of neurons (42 in monkey S and 37 in monkey E), the RDP in the OKN task moved across trials in pseudo-randomized order either in the neuron’s saccade PD or opposite to it. The firing rates of these neurons during the slow-phase were modelled as a function of horizontal and vertical gaze position and velocity (g_x_, v_x,_ g_y,_ and v_y_), as well as the direction of the stimulus motion (MoDir) relative to the neuron’s preferred saccade direction (see Methods, eq. 2). A substantial fraction of neurons exhibited significant sensitivity to gaze position in both horizontal and vertical dimensions (Figure 7, left column). Whereas, consistent with previous reports, a smaller fraction of neurons was sensitive to gaze velocity, with this sensitivity being most pronounced at the beginning of the slow-phase (Figure 7, middle column). Remarkably, most neurons were significantly selective for the direction of stimulus motion that induced fast-phases in their preferred saccade direction, that is, motion opposite to a neuron’s saccade PD. This selectivity was reflected in negative β_MoDir_ values and the significant negative shift of their distribution (Figure 7, right column; Table 3 for summary statistics). The MoDir factor captures a visual motion signal that is independent of the concurrent eye position changes during the OKN slow-phase. Consistent with this interpretation, there was no significant correlation between the MoDir coefficient and the gaze position coefficient along the motion axis (Table 3 for summary statistics). Collectively, these findings indicate that during the OKN slow-phase, LIP neurons encode not only the current position of gaze, but also the ongoing visual motion information associated with the stimulus.

**Table 3.** Wilcoxon signed-rank test on the distribution of β_MoDir_ and its correlation with the gaze position coefficient along the motion axes.

|  |  |  |  | Monkey S |  | Monkey E |  |
| --- | --- | --- | --- | --- | --- | --- | --- |
| | | | | $n = 42$ | | $n = 37$ | |
|  |  |  |  | Beginning of slow-phase | End of slow-phase | Beginning of slow-phase | End of slow-phase |
| Median |  |  |  | -0.06 | -0.1 | -0.08 | -0.09 |
| $p_{\text{distribution}}$ | | | | $4.3 \times e^{-2}$ | $2.2 \times e^{-2}$ | $2.9 \times e^{-2}$ | $2.5 \times e^{-2}$ |
| $p_{\text{correlation}}$ | | | | 0.1 | 0.07 | 0.06 | 0.35 |

**Figure 7.**
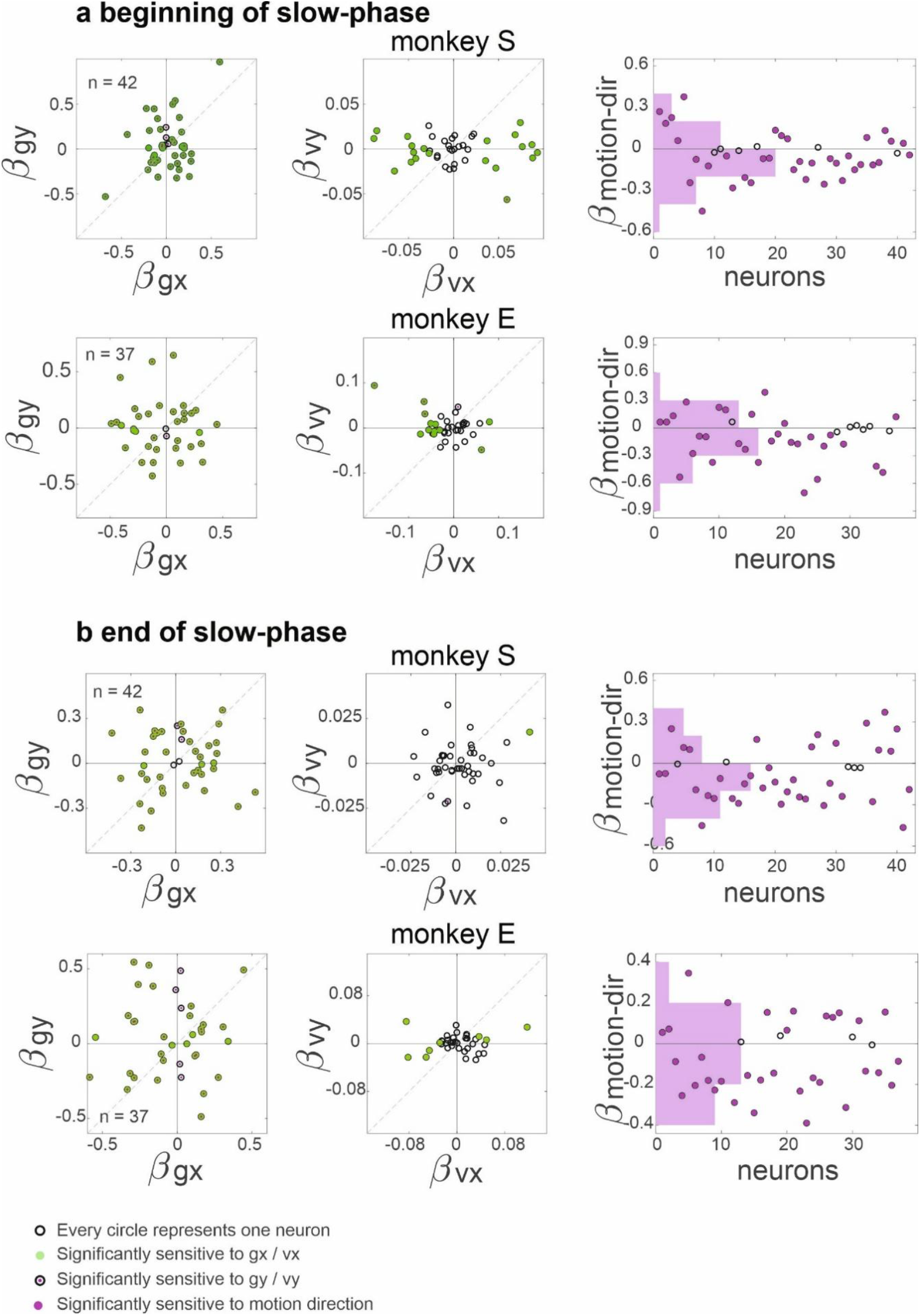
GLM coefficients. Each panel illustrates the GLM coefficients of different factors fitted to the spiking activity of neurons in a 100 ms time window at the beginning (**a**) and at the end (**b**) of the slow-phase of OKN. Every hollow circle represents data from a given neuron. Significant effect of different factors is color-coded as shown in the legend.

### LFP modulations during OKN and saccade

LFPs provide a critical window into the mesoscopic dynamics of cortical processing, capturing the coordinated activity of neuronal populations. In area LIP, much of what we know about eye movement-related activity comes from single-unit recordings, leaving the broader population-level dynamics, especially during reflexive movements, relatively unexplored. Here we compared LFP activity between saccades and OKN fast-phases. Although the two behaviors share similar kinematics, they differ in their underlying control mechanisms: saccades are goal-directed, whereas the fast-phase of an OKN is not voluntarily initiated and lacks explicit motor goals. Given that LFP power reflects local processing demands (44,45) and phase synchrony reflects network coordination (46–48), we used these two measures to compare local and global networks processing in LIP between saccade and OKN fast-phases.

LFP oscillatory activity was analyzed at the beginning and at the end of the OKN slow-phase (after and before the fast-phase, respectively; see Methods), as well as, before and after saccades, using wavelet transformation. LFP power in the low-beta band ([12 20] Hz) was significantly suppressed before and after a saccade (Figure 8a, Table 4 for summary statistics). However, fast-phases were followed by a transient increase in the low beta power at the beginning of the subsequent slow-phase (Figure 8b, Table 4 for summary statistics). In monkey S, this increase also extended to high-beta frequencies ([20-30] Hz, n=38, p=9.8 ×*e*^−4^). Intertrial phase coherence (ITPC, see Methods) in the theta band ([4 8] Hz) increased following both saccade and fast-phases (Figure 9, Table 5 for summary statistics). Different from the findings in delta to alpha band in visual cortex (49,50), as well as theta and beta band in the Frontal Eye Field(49,51), we did not observe a consistent phase-locking between LFP and timing of saccades in LIP. Our findings suggest that, despite differences in local network processing between saccade and OKN fast-phases, LIP engages in at least partly-similar interareal communication after voluntary and reflex fast eye movements.

**Table 4.** Signed-rank test on the average LFP power in a 50 ms time window preceding and following a saccade/fast-phase.

|  | Monkey S | Monkey E |
| --- | --- | --- |
|  | n = 38 | n = 17 |
| Saccade |  |  |
| Testing the power relative to zero |  |  |
| p<sub>pre-saccade</sub> | $2.2 \times e^{-4}$ | $1.9 \times e^{-3}$ |
| p<sub>post-saccade</sub> | $4.6 \times e^{-3}$ | $4.2 \times e^{-4}$ |
| OKN fast-phase |  |  |
| Testing the difference between power preceding and following fast-phase |  |  |
| | $p = 1.6 \times e^{-2}$ | $p = 8.6 \times e^{-3}$ |

**Table 5.** Signed rank test relative to zero on the difference between ITPC in a 50 ms time window preceding and following a saccade/fast-phase.

| Monkey S | Monkey E |
| --- | --- |
| n = 38 | n = 17 |
| Saccade |  |
| $p = 8.8 \times e^{-6}$ | $p = 5.0 \times e^{-4}$ |
| OKN fast-phase |  |
| $p = 1.6 \times e^{-2}$ | $p = 8.6 \times e^{-3}$ |

**Figure 8.**
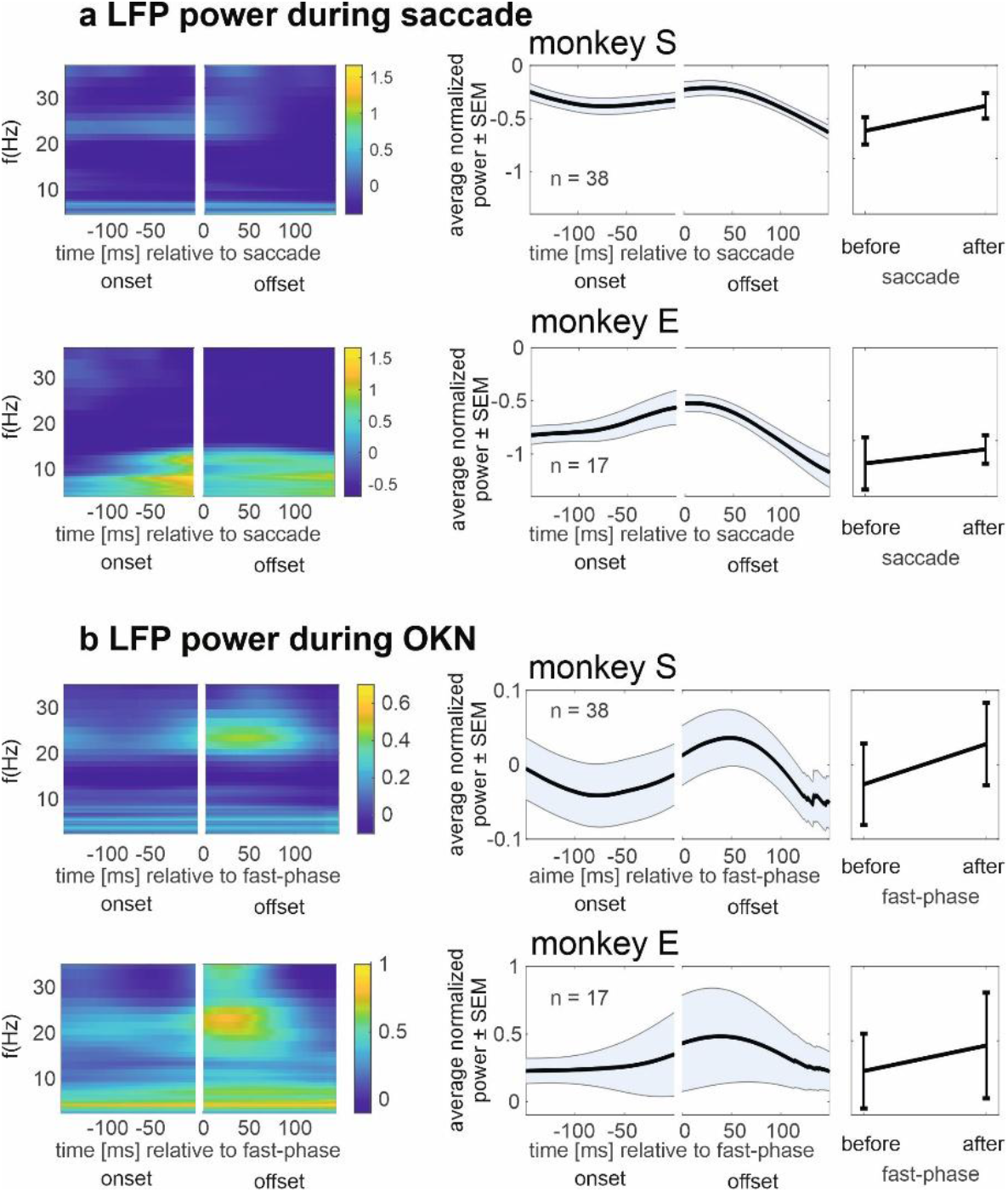
LFP power in the beta band. The left column shows the time-frequency map of LFP power before and after the saccades (**a**) and OKN fast-phases (**b**). The middle column illustrates the time trace of average LFP power in the beta band [12-20] Hz across sessions. The shaded area shows the standard error of the mean across sessions. To plot this figure, the LFP time-frequency power map in every trial was first averaged across frequencies, then the time traces were averaged across all fast-phases/saccades in one session. The right column shows the average power in the beta band in a 50 ms time window preceding the fast-phase/saccade start and in a 50 ms time window following the fast-phase/saccade end. To plot this figure, the LFP time-frequency power map in every trial was first averaged across time and frequencies, then averaged across all fast-phases/saccades in one session.

**Figure 9.**
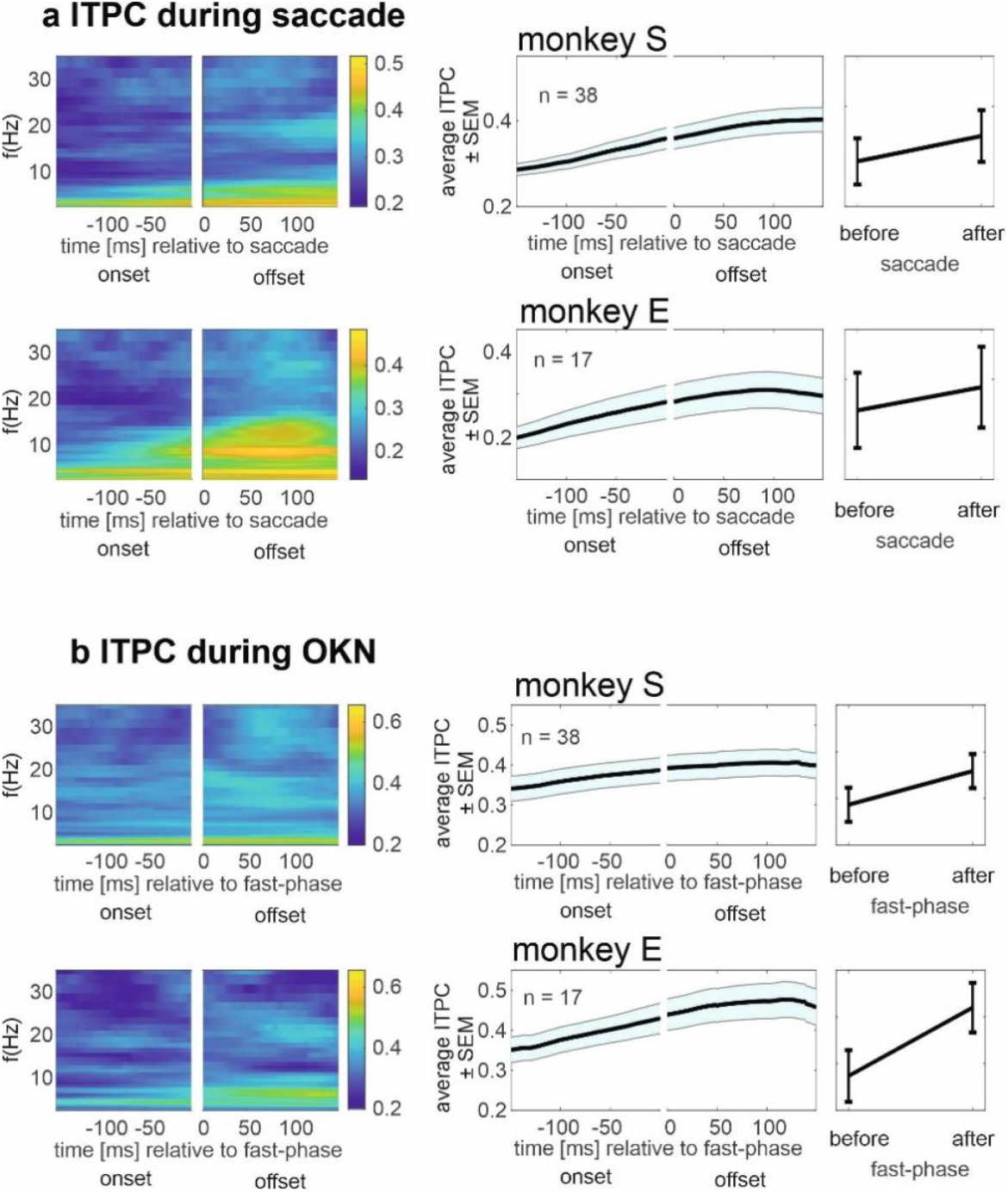
LFP ITPC in the theta band. The left column shows the time-frequency map of LFP intertrial phase coherence (ITPC) before and after the saccades (**a**) and OKN fast-phases (**b**). The middle column illustrates the time trace of the average ITPC in the theta band [4-8] Hz across sessions. The shaded area shows the standard error of the mean. To plot this figure, the ITPC time-frequency map in every trial was first averaged across frequencies, then the time trace was averaged across all fast-phases/saccades in one session. The right column shows the average power in the theta band in a 50 ms time window preceding the fast-phase/saccade start and in a 50 ms time window following the fast-phase/saccade end. To plot this figure, the ITPC time-frequency map in every trial was first averaged across time and frequencies, then averaged across all fast-phases/saccades in one session.

## Discussion

In this study we have compared neural activity in macaque area LIP across voluntary, visually guided saccades, and reflexive OKN fast phases with similar metric (direction and amplitude). For most neurons, spiking activity across a fast phase was attenuated or even absent as compared to its peri-saccadic activity. While beta-band power differentiated saccades from OKN fast-phases, theta-band phase coherence increased after both movements, suggesting distinct local processing but overlapping interareal coordination.

### Distinct role for the control of fast eye-movements in area LIP

The differences we observed in the main-sequence relationships between saccades and OKN fast-phases provide strong evidence that these two eye movements, despite their similar kinematics, are controlled by partially distinct oculomotor circuits. Specifically, OKN fast-phases exhibited significantly lower peak velocities than saccades of comparable amplitude, consistent with earlier behavioral observations in both humans and non-human primates (20,30,52). Such differences are generally interpreted as reflecting divergent upstream control mechanisms rather than differences at the level of the brainstem burst generators, which are thought to be largely shared (53).

Voluntary saccades rely heavily on cortical structures involved in goal selection and decision-making, including area LIP and the frontal eye fields (FEF) (40,54). In contrast, OKN fast-phases are reflexive resetting movements embedded within a continuous sensorimotor loop driven primarily by visual motion signals and brainstem–cerebellar circuitry (55). The altered main-sequence therefore aligns well with the idea that cortical contributions to OKN fast-phases are weaker or qualitatively different from those involved in voluntary saccades, a view in line also with the neural data reported here and discussed below.

### Reduced LIP activity during OKN fast-phases

The reduced spiking activity in LIP during OKN fast-phases compared to saccades of matched amplitude and direction parallels the behavioral dissociation suggested by the main-sequence analysis. Neurons that were robustly tuned for saccade direction showed strong perisaccadic modulation but little or no modulation around fast-phase onset. This pattern was consistent across single neurons and at the population level and was evident both before and after movement onset. These findings are well aligned with the view that area LIP is more strongly engaged in the planning, evaluation, and monitoring of voluntary eye movements than in their direct execution (2,40,56). Numerous studies have demonstrated that LIP neurons encode saccade goals, expected outcomes, and decision variables rather than purely motor commands (54,57,58). Bremmer and colleagues further showed that LIP activity reflects spatial goals and integrates visual, eye position, and motor-related signals relevant for voluntary action (2,3,37,38,59). In this framework, the weak engagement of LIP during OKN fast-phases is expected: fast-phases lack explicit motor goals, are not selected among alternatives, and do not require monitoring against an intended target. Our findings strengthen the interpretation that LIP activity during saccades reflects higher-level control and monitoring processes rather than a generic response to eye movement kinematics.

### Absence of modulation by fast-phase frequency or amplitude

Functional imaging studies in humans had reported systematic changes in parietal BOLD activity as a function of fast-phase frequency, distinguishing stare nystagmus from look nystagmus (22,35). In contrast, we found no evidence that either the frequency or amplitude of OKN fast-phases modulated spiking activity in macaque area LIP. This absence of modulation further supports the conclusion that LIP is not involved in monitoring fast-phase execution parameters. If area LIP was encoding aspects of fast-phase generation or evaluation, one might expect activity to scale with movement frequency or size, as observed for voluntary saccades (1–3). Instead, the insensitivity to fast-phase statistics suggests that LIP largely ignores these reflexive movements once they are triggered, consistent with their limited behavioral relevance.

The discrepancy with human fMRI findings likely reflects differences in what the two signals capture. BOLD responses integrate synaptic and modulatory activity over extended periods of time and may reflect changes in global network state or attentional engagement rather than fast-phase-specific computations (23). In contrast, single-unit spiking in LIP appears tightly linked to voluntary control demands, which are minimal during OKN.

### Gaze-related signals and unexpected sensitivity to visual motion during slow-phase

Our GLM analysis during OKN slow-phases revealed robust encoding of eye position in both horizontal and vertical dimensions, fully consistent with previous work (36–38,59,60). These signals are thought to provide a substrate for transforming visual information into motor and spatial representations across reference frames (12), and their persistence during involuntary behavior underscores their fundamental role in parietal cortex function. More surprising was the modulation of LIP activity by the direction of visual motion driving the OKN, independent of eye position and velocity. LIP responses to visual motion have been reported previously, even when motion was behaviorally irrelevant (39,40,43), and (7) showed that some LIP neurons are selective for pursuit direction. However, the robust motion-direction sensitivity observed here during OKN slow-phases suggests that LIP may monitor salient motion in the external world even during reflexive behavior. One possible interpretation is that area LIP continues to encode task-relevant environmental dynamics that could inform higher-level perceptual or behavioral decisions, even when the eye movement itself is not voluntarily controlled. This interpretation aligns with more recent views of LIP as part of a broader perceptual - cognitive network that tracks both internal state (e.g., gaze direction) and external sensory structure (41).

### LFP dynamics: local processing versus network-level coordination

Our results on LFP dynamics provide an important mesoscopic perspective that complements the spiking data. Beta-band power was suppressed around saccades but transiently increased following OKN fast-phases, suggesting distinct local processing demands for voluntary versus reflexive movements. Beta suppression has been widely associated with motor preparation and active sensorimotor processing (61), consistent with the engagement of LIP in voluntary saccades. In contrast, the shared increase in theta-band phase coherence following both saccades and fast-phases suggests that LIP participates in similar large-scale network coordination after fast eye movements, regardless of their voluntary nature. Theta-band synchronization has been linked to long-range communication and cognitive control across cortical areas (62), and similar post-movement coordination may support perceptual updating after any rapid gaze shift. The dissociation between power and phase measures therefore mirrors the results of spiking neural activity: local processing in area LIP differentiates strongly between voluntary and reflexive movements, while broader interareal coordination appears at least partly shared.

### Conclusions

Taken together, our findings support a view in which area LIP plays a prominent role in the monitoring and cognitive control of voluntary saccades but is only weakly engaged during reflexive fast phases. While area LIP continues to encode gaze position and visual motion during involuntary behavior, it appears largely insensitive to fast-phase execution parameters themselves. This functional specialization is reflected consistently across behavioral dynamics, spiking activity, GLM-based encoding, and LFP oscillations, reinforcing the view of area LIP as a key area for integrating sensory and cognitive signals relevant for goal-directed action rather than a generic oculomotor controller.

## Acknowledgements

This work was supported by Deutsche Forschungsgemeinschaft (CRC/TRR-135 – Cardinal Mechanisms of Perception, project No. 222641018, and EXC 3066/1 The Adaptive Mind, project No. 533717223), and the European Union (PLACES).

